# Three dimensionally printed microstructured alginate scaffolds for neural tissue engineering

**DOI:** 10.1101/2024.04.23.590678

**Authors:** Jianfeng Li, Benjamin Hietel, Michael G.K. Brunk, Armin Reimers, Christian Willems, Thomas Groth, Holger Cynis, Rainer Adelung, Fabian Schütt, Wesley D. Sacher, Joyce K.S. Poon

## Abstract

The integration of scaffolds, signalling cues, and cellular components is essential in tissue engineering to create an *in vivo* equivalent environment that supports physiological function. Scaffolds provide mechanical reinforcement for cellular proliferation and differentiation while providing cues that instruct the development of cells during culture. Alginate (Alg) is a versatile biopolymer for scaffold engineering. However, due to a lack of intrinsic cell-binding sites, thus far, Alg must be functionalized for cellular adhesion. Here, we demonstrate proof-of-concept, bioactive additive-free, microstructured Alg (M-Alg) scaffolds for neuron culture. The M-Alg scaffold was formed by introducing tetrapod-shaped ZnO (t-ZnO) microparticles as structural templates in the Alg that were subsequently removed. These transparent, porous, additive-free Alg-based scaffolds with neuron affinity are promising for neuroregenerative and organoid-related research.

**Highlights:** Tetrapod-shaped ZnO (t-ZnO) microparticles are used as a template for the fabrication of open interconnected channels and textured surfaces in 3D printed microstructured alginate (M-Alg) scaffolds.

Primary mouse cortical neurons seeded on the 3D printed M-Alg scaffolds show improved adhesion and maturation with extensive neural projections forming inside the scaffolds.

## 1. Introduction

*In vitro* three-dimensional (3D) cellular models are a method for replicating the physiological microenvironments of *in vivo* tissues, serving as useful supplements to animal models and *ex vivo* tissues. These 3D models complement pathophysiological models and elucidate the mechanisms that govern cellular behaviours for advancing our understanding of various diseases [1]. 3D neural models serve as an analogue of their *in vivo* counterpart for investigating therapies for neurological disorders and studying the mechanisms of neural computation. Among the various fabrication methods for 3D cell models, 3D bioprinting stands out for its ability to fabricate models with precise anatomical features and to incorporate multiple bioactive components in a programmable fashion [2, 3]. Although extrusion-based 3D bioprinting, which utilizes bioink to encapsulate living cells, is popular, the effective disinfection of cell-laden ink and mitigation of shear stress on cell viability during the printing process are critical challenges [4, 5]. An alternative approach is to initially print a scaffold, followed by sterilization and cell seeding [6, 7]. However, usually, cells are confined to the surface of the scaffold, and the internal volume remains devoid of cells owing to the barrier properties of the printed material. This may result in diminished cell-loading capacity and compromised intercellular communication.

Alginate (Alg), which has a similar linear polysaccharide structure to the naturally-occurring hyaluronic acid in the brain extracellular matrix, is a well-established biopolymer derived from brown algae and has found numerous applications since its discovery over a century ago [8, 9]. In particular, Alg solutions can form gels through ionic crosslinking in a biocompatible manner [9], eliminating the need for additional chemical modifications [10] or the use of toxic agents such as carbodiimide or other chemical cross-linkers [11]. Alg has been extensively applied in 3D bioprinting since it is easy to print and biocompatible [12, 13]. Cells encapsulated in the Alg have superior performance in 3D printed Alg compared to bulk Alg hydrogels devoid of printed channels because of the improved transport of nutrients and oxygen to the cells [14]. Because Alg lacks cell-anchoring ligands, cell seeding on Alg scaffolds does not work. Thus, in the printing process, cells are encapsulated in the Alg-containing ink, but printing such cell-laden inks applies shear stress to the cells and raises the risk of contamination, reducing the viability of printed cells [15]. Modifying the Alg ink or the surface of Alg scaffolds with bioactive components (e.g., arginine-guanidine-aspartate and collagen) or functional coatings can promote cell attachment and growth [6, 9, 16], but may not always be compatible with the intended chemical or physiological tests using the scaffold.

In this study, we present a technique for creating transparent, additive-free, microstructured Alg (M-Alg) scaffolds with varying diameter channels and textured surfaces that have high cellular affinity as illustrated in **Figure 1**. Using tetrapod-shaped ZnO (t-ZnO) microparticles as templates, which are later removed, we produced scaffolds that encouraged neuron adhesion, proliferation, and circuit formation without the need for pre-treatments or bioink modifications. Interconnected channels in scaffolds offer pathways for vascularization, nutrient transport, waste disposal, and cell migration. Multi-sized channels are particularly beneficial for tissue regeneration [6, 17]. Alg scaffolds with uniformly sized channels have previously been shown to promote blood vessel growth without inflammation or abnormalities after three weeks of implantation [18] and facilitate controlled delivery of therapeutics [19]. Compared to pristine Alg-based (P-Alg) scaffolds, in a series of proof-of-concept experiments, the M-Alg with the interconnected channels were found to be more conducive to the adhesion, proliferation, migration, and establishment of active circuits of primary mouse neurons seeded on the structure. We hypothesize that the internal channels and textured surfaces [20] enhanced the efficacy of M-Alg scaffolds in peripheral nerve generation based on the previously reported results [21]. This approach of forming porous Alg scaffolds reduces variability in cellular responses, offering a potential alternative to Matrigel as a biomaterial.

**Figure 1.**
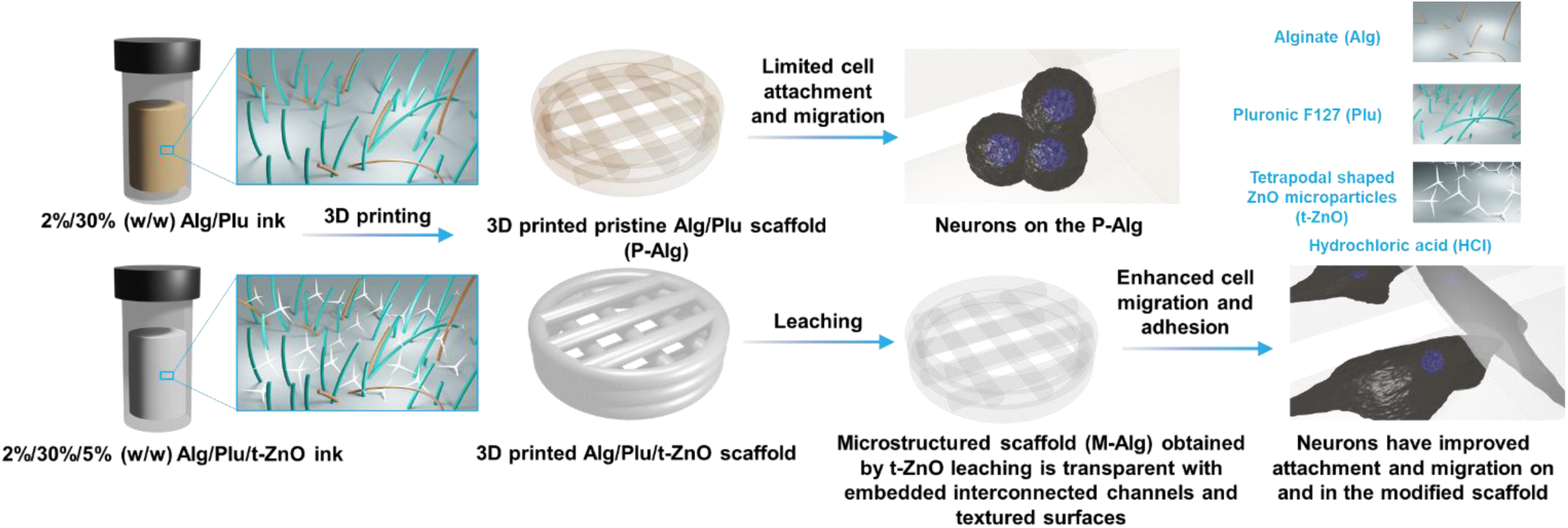
Schematic depiction of the 3D printing process contrasting pristine Alg based scaffold (P-Alg) and microstructured Alg scaffold (M-Alg) with interconnected channels and textured surfaces. Neurons exhibit improved adhesion and proliferation on the M- Alg, while showing limited adhesion on the P-Alg.

## 2. Materials and Methods

### 2.1. Ink preparation and 3D printing

Alg with an average molecular weight (Mn) of 80,000-120,000 from brown algae and Pluronic F-127 (Plu) with an Mn of 12,600 were purchased from Merck (Germany).

For 2%/30% (w/w) Alg/Plu ink preparation, 200 mg Alg and 3,000 mg Plu were mixed mechanically with 10.000 g deionized water (DI water) at 4 °C for 12 h. Using the prepared 2%/30% (w/w) Alg/Plu ink, 2%/30%/5% (w/w) Alg/Plu/t-ZnO ink was prepared by adding 500 mg t-ZnO into the solution and mixed mechanically at 4 °C for another 12 h.

The as-prepared inks were loaded into a 3D printing syringe barrel (Nordson EFD, United States) and centrifuged to de-bubble at 1,000 xg for 10 min at 4 °C (Fisherbrand Centrifuge, Germany). 3D printing was performed using an extrusion-based 3D printer (RegenHU, Switzerland) with a G20 needle (Nordson EFD, United States) fitted in a syringe barrel at room temperature (RT, 21 °C). An ink feeding speed of 6 mm/s was used, with the corresponding applied air pressure for each ink (2%/30% (w/w) Alg/Plu ink: 130 kPa; 2%/30%/5% (w/w) Alg/Plu/t-ZnO ink: 170 kPa), followed by 10 min of crosslinking in a 2% (w/v) calcium chloride (CaCl_2_; Merck, Germany) solution at RT. Thereafter, 37% hydrochloric acid (HCl; Fisher Chemical, Germany) was used to remove the t-ZnO component from the crosslinked scaffolds for 3 min, and the scaffolds were reinforced in 2% (w/v) CaCl_2_ for another 5 min. The obtained microstructured scaffolds were immersed and washed with copious amounts of DI water to remove the acid residues from the previous step. Before being seeded with cells, the 3D printed scaffolds underwent a sterilization process by immersing in 70% (v/v) ethanol solution for 20 min, followed by desiccation in a biosafety cabinet (BSC). A subsequent sterilization procedure was carried out under UV irradiation for 1 h. The sterilized scaffolds were immersed in culture medium overnight in a cell culture incubator (37 °C with humidified 5% CO_2_; Binder, Germany) before cell seeding. The t-ZnO microparticles used here were synthesized via a previously reported flame transport method [22].

### 2.2. Primary cortical neuron isolation and culture

Primary mouse cortical neurons were isolated from the cortices of mouse (strain C57BL6/J) embryos (E17) as reported previously [23]. The primary cortical cell culture medium comprised Neurobasal^TM^ Plus Medium (Gibco, USA) supplemented with 2% (v/v) B27^TM^ Plus Supplement (Gibco, USA), 1% (v/v) GlutaMAX™ Supplement (Gibco, USA), and 1% (v/v) penicillin-streptomycin (10,000 U/mL; Gibco, USA). The initial cell seeding density for the 3D scaffolds was 0.8×10^5^ cells/cm^2^. Cells were cultured in a cell culture incubator, and one-third of the culture medium was changed every three days.

### 2.3. Scanning electron microscopy

For morphology study of 3D printed scaffolds without cells, samples were imaged using a Hitachi TM4000Plus scanning electron microscope (SEM) (Hitachi, Japan) equipped with an ultracool stage at -30 °C (Deben, UK). For scaffolds with cells, samples were pretreated by immersing and fixing in 4% paraformaldehyde solution (PFA; ThermoFisher Scientific, Germany) for 30 min. After washing three times with a 2% (w/v) CaCl_2_ solution, the samples were imaged using an SEM equipped with an ultracool stage at -30 °C. Cross-sectional imaging was performed by fracturing samples using a surgical scalpel.

### 2.4. Cell Viability Evaluation

For cell viability assessment, staining was performed using 8 μM Calcein AM (CA; Merck, Germany) for live cell staining, 2 μM propidium iodide (PI; Merck, Germany) for dead cell staining, and 20 μM Hoechst 33342 (HO; Miltenyi Biotec, Germany) for visualization of nuclei. Briefly, cell-containing structures were incubated with CA for 15 min in a cell culture incubator, followed by addition of PI and HO to the staining medium for another 15 min of incubation. After washing the stained structures with fresh medium, a confocal microscope (Zeiss LSM 900, Germany) was used for imaging.

### 2.5. Proliferation

The proliferation of neurons on different 3D scaffolds was assessed using PrestoBlue^TM^ cell viability reagent (ThermoFisher Scientific, Germany) according to the manufacturer’s instructions. Briefly, cell-seeded scaffolds were incubated with 500 µL 10% (v/v) reagent culture medium solution for 30 min in an incubator on days 1 and 7. Subsequently, the supernatant was transferred to a 96-well plate for fluorescence intensity measurement using a microplate reader (excitation/emission: 544/590; POLARstar Omega, Germany). The fluorescence value of the cell-seeded P-Alg samples on day 1 was set to 100, and the proliferation on the other samples on days 1 and 7 was calculated relative to this baseline.

### 2.6. Calcium imaging

Neuronal activity was assessed by staining the neurons with 5 μM Calbryte 520 AM (AAT Bioquest, USA) and 0.04% Plu for 30 min in a cell culture incubator. Plu was used to enhance the cell loading and staining performance of the Calbryte 520 AM. After washing the stained structures with fresh medium, a confocal microscope (Zeiss LSM 900, Germany) with a mounted cell culture incubator (XLmulti S2 DARK, Germany) was used for imaging.

### 2.7. Immunostaining

Phosphate-buffered saline (PBS; diluted from 10× PBS pH 7.4, ThermoFisher Scientific, Germany) and 4% PFA solution were supplemented with 5 mM CaCl_2_ to prevent the dissolution of the crosslinked Alg structure. The cell-seeded scaffolds were removed from the culture medium and washed 3 times with PBS. Subsequently, the cells were fixed with 4% PFA for 30 min at RT. After rinsing three times with PBS, the cells were permeabilized and blocked with 0.3% (v/v) TritonX-100 (Calbiochem, Germany) and 10% (v/v) horse serum (HS; MP Biomedicals, Germany) in PBS for 2 h at RT. Subsequently, the scaffold was incubated overnight at 4 °C with anti-β-tubulin III antibody (1:100; Merck, Germany) and anti-synaptophysin antibody (1:100; Merck, Germany) in a 10% (v/v) HS PBS solution. The scaffold was then rinsed three times with PBS and incubated with Alexa Fluor™ 555 conjugated secondary goat anti-mouse IgG antibody (1:100; Merck, Germany) and CF™ 633 conjugated secondary donkey anti-rat IgG antibody (1:100; Merck, Germany) for 2 h at RT, followed by incubation with AlexaFluor™ 488 phalloidin (2 drops/mL; ThermoFisher Scientific, Germany) and 4’,6-diamidino-2-phenylindole (DAPI, 2 μg/mL; Merck, Germany) for 1 h at RT. Finally, the scaffold was washed with PBS for 3 times. Images were acquired using a Zeiss LSM 900 confocal microscope.

### 2.8. Statistical analysis

The proliferation test measurements were conducted in quintuplicate, and the data are presented as the mean ± standard deviation. Data plotting and statistical analyses were performed using Origin 2022b (OriginLab, USA). Since the Brown-Forsythe test for data variance homogeneity was not satisfied (P < 0.05), a significance level of 0.01 was employed for the two-way ANOVA (Bonferroni post hoc test).

## 3. Results and Discussion

### 3.1. 3D printed Alg-based scaffold with open interconnected channels and textured surfaces

**Figure 2.**
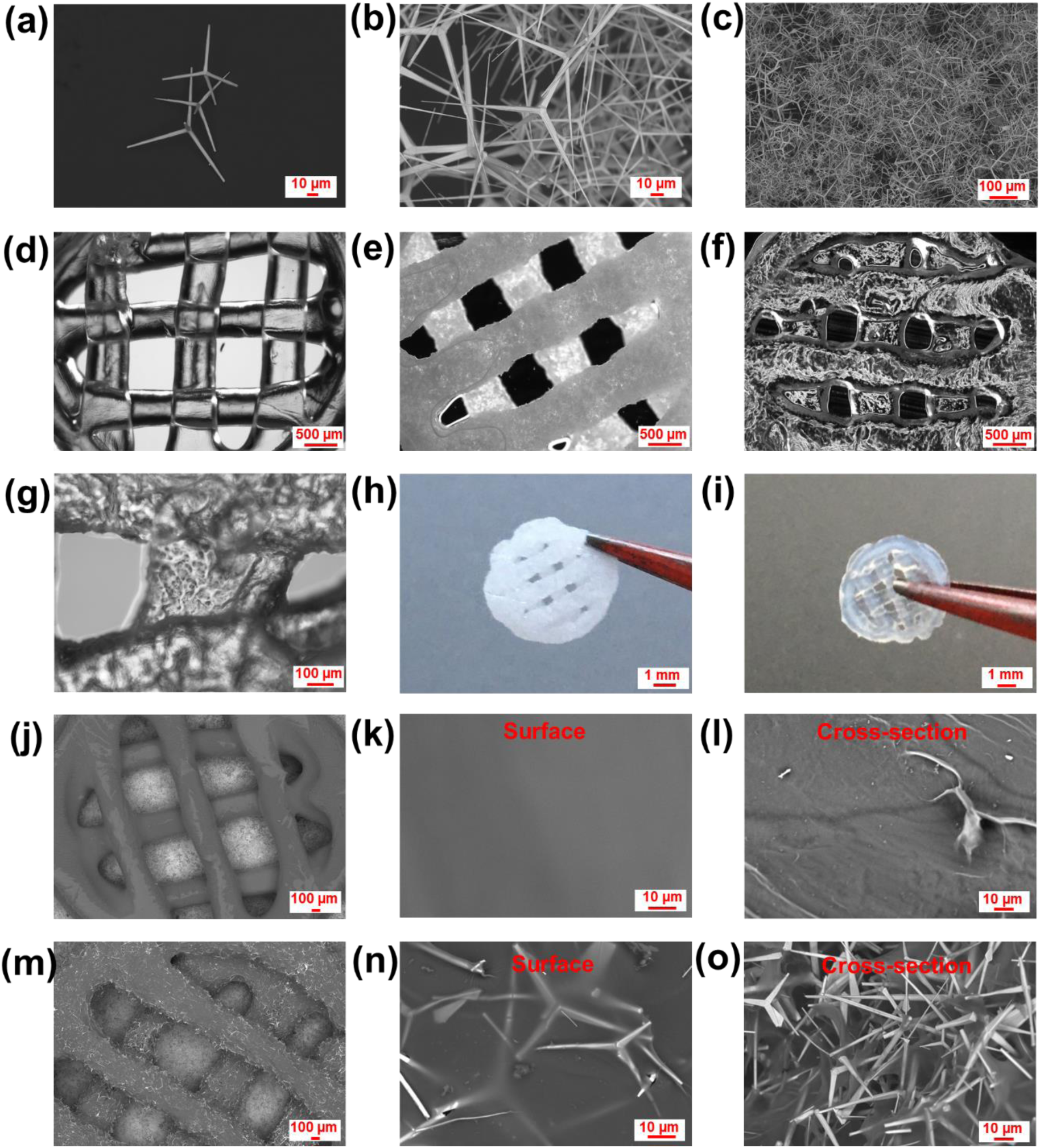
Morphology of 3D printed P-Alg and M-Alg scaffolds. (a-c) SEM images of different concentrations of t-ZnO microparticles. Optical microscope images of 3D printed (d) P-Alg scaffold, (e) 2%/30%/5% (w/w) Alg/Plu/t-ZnO scaffold, and (f-g) M- Alg scaffold at various magnifications. Photographs of (h) 3D printed 2%/30%/5% (w/w) Alg/Plu/t-ZnO scaffold and (i) M-Alg scaffold. SEM images of an overview of (j) 3D printed P-Alg scaffold with magnified (k) surface and (l) cross-sectional morphology. SEM images of (m) 3D printed 2%/30%/5% (w/w) Alg/Plu/t-ZnO scaffold with magnified (n) surface and (o) cross-sectional morphology.

Alg is widely recognised for its versatility in biomedical applications, but its use in tissue engineering has been limited by the absence of bioactive ligands. Enhancing the cellular affinity of Alg without functionalization could expand its use by reducing complications from foreign materials. We use tetrapod-shaped ZnO (t-ZnO) microparticles as templates to modify the morphology and internal structure of 3D-printed Alg scaffolds. The t-ZnO microparticles, characterized by four arms, naturally assemble into interlinked networks in close proximity (**Figure 2a-c**). Remarkably, these microparticles can be easily eliminated using a hydrophilic volatile acid (such as HCl), ensuring the absence of toxic residues post-fabrication, which is preferred over conventional particulate leaching using organic solvents, such as dimethylformamide [24]. The acid treatment does not affect the morphology and surface of 3D printed Alg scaffolds because the Alg structure is chemically inert to HCl [25].

The 3D-printed Alg scaffold exhibited optical transparency (**Figure 2d**). However, upon integrating t-ZnO microparticles, the structure assumed a white hue but the print quality was preserved (**Figure 2e, h, and Figure S1a**). Our previous study indicated that inks containing t-ZnO microparticles exhibit favourable rheological attributes at RT for 3D printing [13]. Leaching in HCl reinstates the transparency of the structure, along with a textured morphology, while preserving mechanical integrity (**Figure 2f, g, i, Figure S1b,** and **Video S1**). Incorporating t-ZnO microparticles into Alg-based ink yields a roughened printed framework, which persists as a network within the hydrogel matrix (**Figure 2j-o**).

### 3.2. Neuron adhesion, viability and proliferation on 3D scaffolds

**Figure 3.**
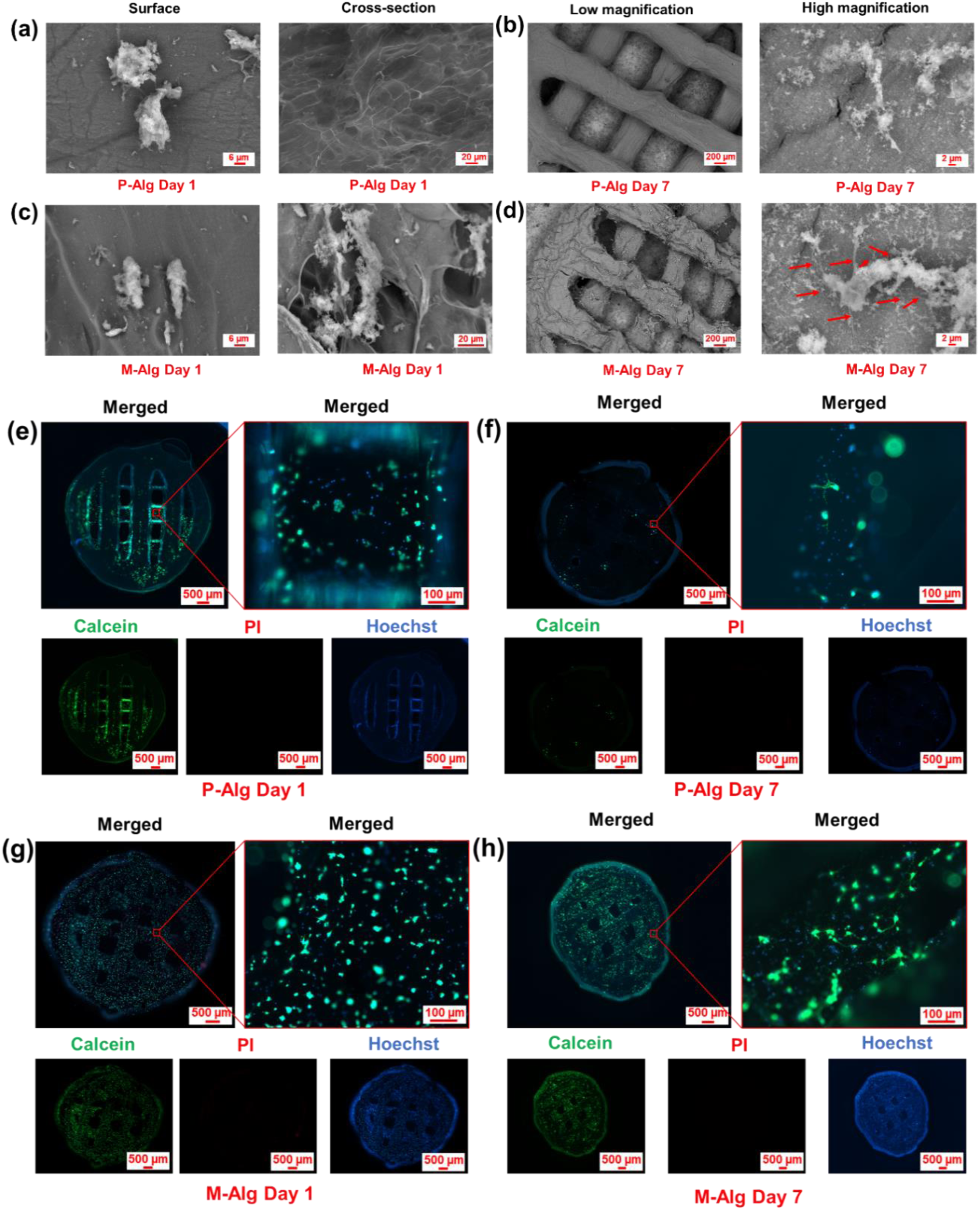
Adhesion and survival of primary cortical neurons on the P-Alg and M-Alg scaffolds. SEM images highlighting neurons on the surface and interior of the (a) P-Alg and (c) M-Alg scaffolds on Day 1. SEM images showing the scaffold and neuron morphology of (b) P-Alg and (d) M-Alg at different magnifications on Day 7, with red arrows indicating neurites. Fluorescence microscope images of cells on P-Alg on (e) Day 1 and (f) Day 7, and on M-Alg on (g) Day 1 and (h) Day 7.

Cell adhesion and viability within supportive structures are crucial to successful engineering of tissues, affecting cellular migration, proliferation, and differentiation [26]. Primary cortical neurons adhered initially to both P-Alg and M-Alg scaffolds; however, the M-Alg scaffold demonstrated significantly better and more uniformly distributed cellular adhesion with neurons that exhibited a spread morphology (**Figure 3a, c, e, and g**). Neurons seeded onto the M-Alg scaffolds could infiltrate the open channels, substantially increasing the filling of the scaffold architecture (**Figure 3c**). Despite the initial cellular adhesion on both scaffold types, adhesion on P-Alg scaffolds was only temporary with a round morphology of cells followed by a markedly decrease in cellular presence by Day 7 (**Figure 3b, d, f, and h**). In contrast, neurons formed durable connections on M-Alg scaffolds, extending neurite anchors throughout (**Figure 3d**) and maintained high-density populations with complex neural networks by Day 7 (**Figure 3h**). After a 7-day culture period, the P-Alg scaffolds appeared smooth, whereas M- Alg scaffolds showed a rough texture with visible t-ZnO shapes, due to internally connected channels (**Figure S2a-d**). These results show the benefit of textured, rough scaffolds based on M-Alg with internal channels promoting neuron adhesion and growth in contrast to the smooth and hydrophilic characters of P-Alg scaffolds, which obviously did not support durable attachment of cells.

**Figure 4.**
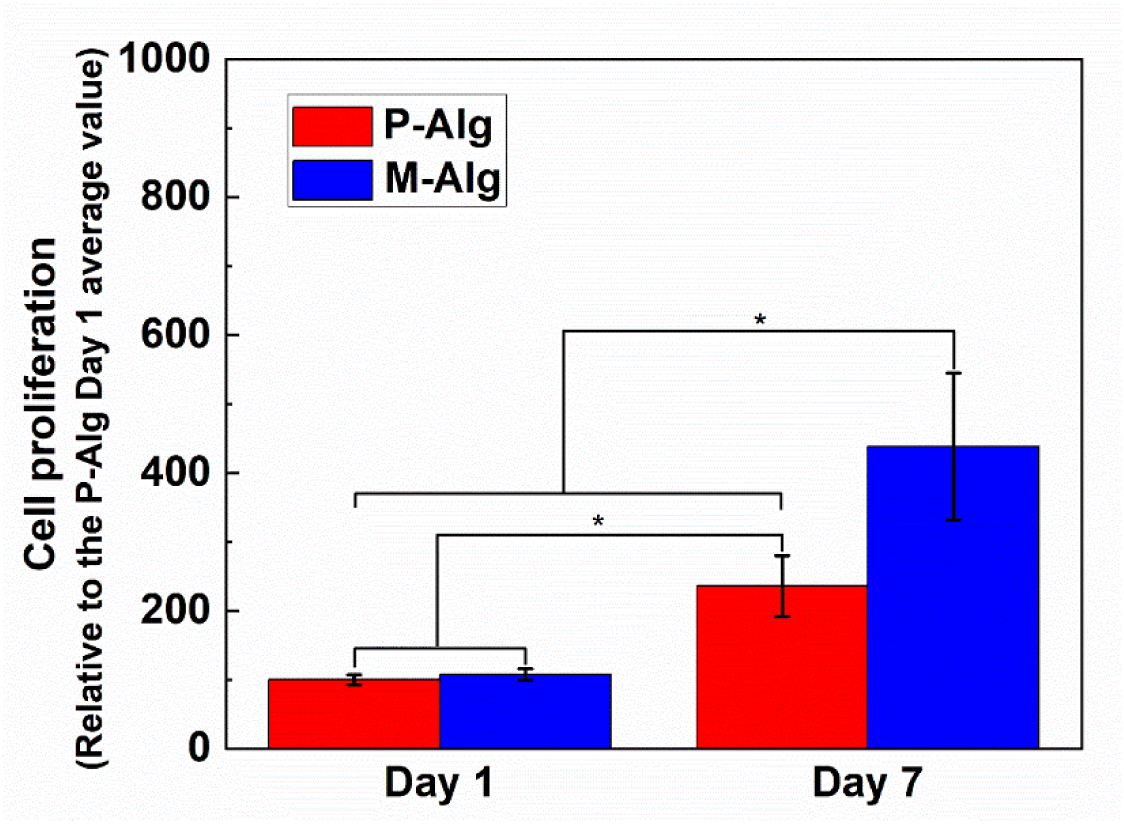
Neuron proliferation on both P-Alg and M-Alg scaffolds by relative fluorescence intensity increase compared to the P-Alg Day 1 average value. Mean ± SD, n = 5. [Bonferroni post hoc, P < 0.01 (M-Alg Day 7 vs all comparisons; P-Alg Day 7 vs P-Alg Day 1 and M-Alg Day 1)]

The influence of P-Alg and M-Alg scaffolds on neuronal proliferation was tested with the PrestoBlue assay. As shown in **Figure 4**, neurons exhibited proliferation on both scaffolds, with the number of neurons on M-Alg significantly higher than the P-Alg groups. The significantly higher neuron numbers observed on M-Alg Day7 may be attributed to the increased surface area conductive to neuron growth and enhanced nutrition/oxygen transport facilitated by the embedded channels within the scaffold. Additionally, the softer nature of the porous scaffold based on M-Alg could potentially contribute to the enhanced neuron growth [27]. Two-way ANOVA analyses revealed the significant influence of both scaffold type [F (1, 36) = 29.8, P < 0.01] and neuron culture time [F (1, 36) = 147.2, P < 0.01] on neuron proliferation. Thus, M-Alg scaffolds hold promise as improved substrate supporting neuronal cell proliferation.

### 3.2. Neuron maturation evaluation

**Figure 5.**
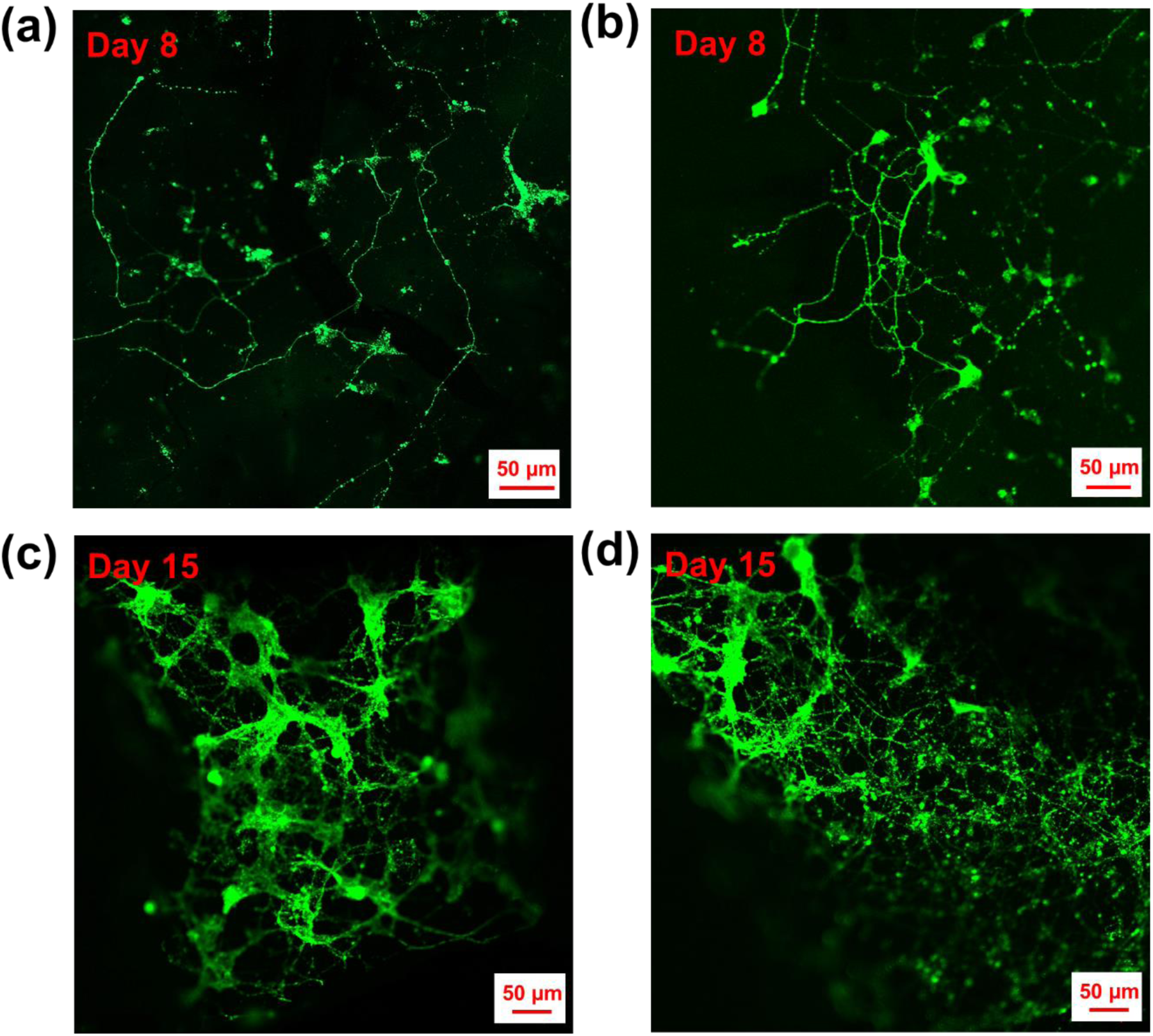
Fluorescence microscopy images showing extended neuronal networks formed in the M-Alg scaffolds on Day 8 and Day 15. Neuronal network staining was done with Calbryte 520 (green).

Neuronal cells develop morphological, electrophysiological, and molecular features as they mature [28]. To evaluate neuron maturation with the M-Alg scaffold, 3D neuronal cultures were stained with calcium-sensitive dye Calbryte 520 and markers of β-tubulin III and synaptophysin to differentiate neuronal phenotypes. Significant neuron outgrowth, spanning several hundred micrometres, was observed in the M-Alg scaffolds on Day 8, as illustrated in **Figure 5a-b** after staining with Calbryte 520 indicating also functional activity by presence of intracellular free calcium [29]. With an additional week of culture, an even more complex assembly of neuronal networks emerged, accompanied by a widespread distribution of dendritic spines throughout the network (**Figure 5c-d**). This development suggests the potential for increased plasticity in these neuronal networks when cultured in the M-Alg scaffolds [30].

**Figure 6.**
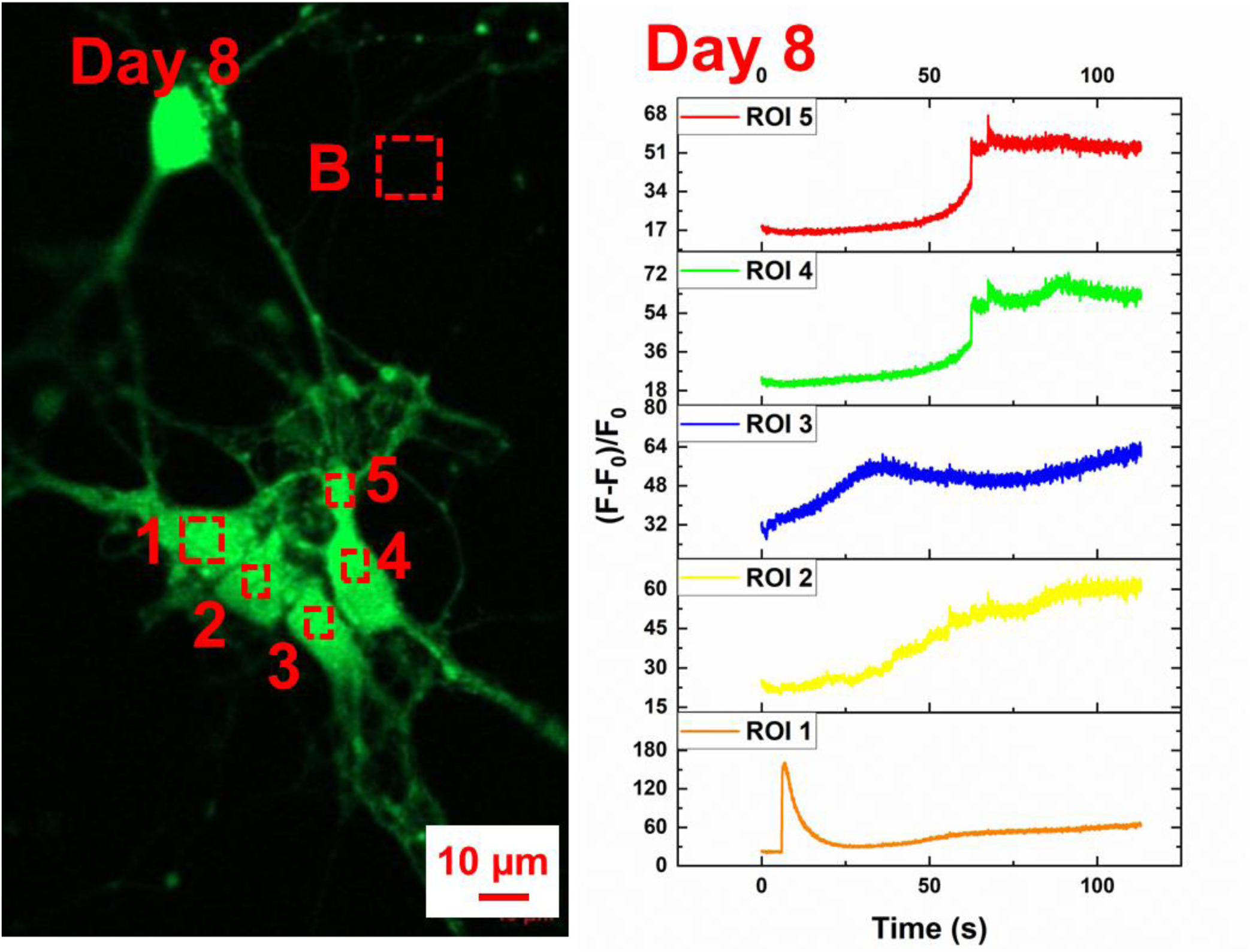
The neuronal networks formed in the M-Alg scaffold show spontaneous activity on day 8, indicating active intercellular connections formed within the scaffold. Neuronal network stained by Calbryte 520 (green). The numbers (1-5) specify five regions of interest (ROIs) and the letter B specifies a background area for subtraction during fluorescence signal processing.

Spontaneous neuronal activity was detected in the established neuronal network formed in the modified M-Alg scaffold by Day 8 (**Figure 6 and Video S2**). This activity revealed signal transmission among cells, indicating that the complex intercellular connections that had been formed were functioning.

**Figure 7.**
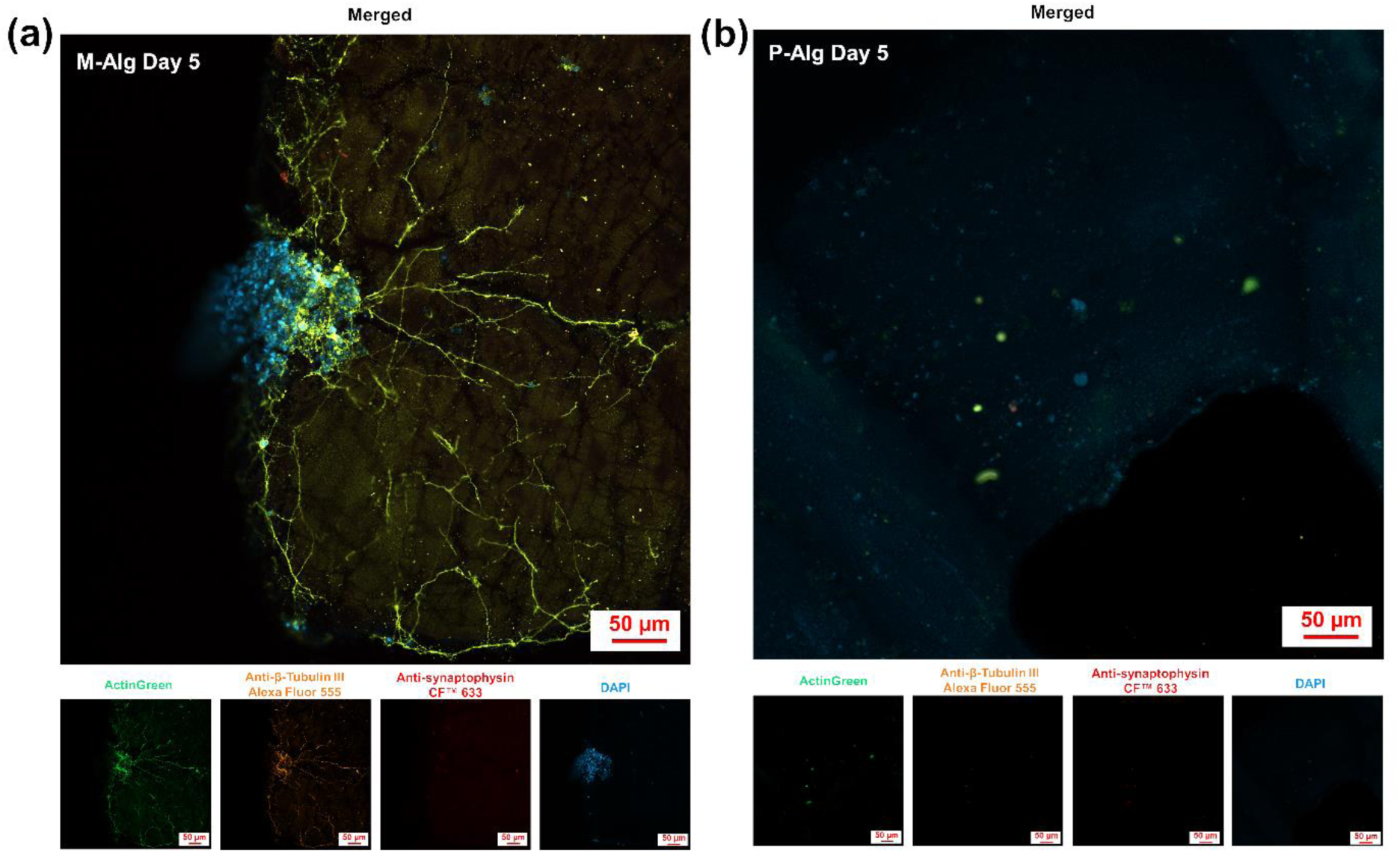
Neuronal maturation marker expression in different 3D cultures. Immunofluorescence microscopy of neurons cultured on (a)M-Alg and (b) P-Alg on Day 5. Robust neural network development observed in M-Alg scaffold compared to sparse neurons on P-Alg scaffold by Day 5. Actin (ActinGreen; green), β-tubulin-III (Alexa Fluor 555; orange), synaptophysin (CFTM 633; red) and nuclei (DAPI; blue).

Consistent with the calcium imaging in **Figure 5**, actin-staining showed densely branched neuronal networks in the M-Alg scaffold on Day 5 (**Figure 7a and Figure S3**). This stands in stark contrast to the notably sparse distribution of neurons observed on the P-Alg scaffold on the same day, as well as within both scaffolds on Day 1 (**Figure 7b** and **Figure S4a-b**). Additionally, staining for the neuronal differentiation marker β-tubulin-III confirmed the development of a complex neuronal network within the M-Alg scaffold, as shown in **Figures 7a-b and S3**. The presence of synapses, as evidenced by synaptophysin staining, indicates synaptic vesicle formation along the neurites exclusively in the M-Alg scaffold, showing its capability to support complex neuronal network development and synaptic connectivity, which can be considered as further evidence for the functionality of neuronal networks supported by the porosity and roughness of these type of scaffolds (**Figure 7a-b and Figure S3**).

## 4. Conclusion

In summary, we have described a simple fabrication strategy to create in-strand open interconnected channels and textured surfaces in the Alg-based scaffold by combination with t-ZnO crystals as template structure that can be easily dissolved after printing. In contrast to the conventional ways of introducing external bioactive moieties or molecules, such as conjugating arginine-glycine-aspartic acid (Arg-Gly-Asp, RGD) [31] or mixing with biomaterials (collagen [9], graphene oxide [12]), to increase the bioactivity of 3D Alg structures, our approach physically modifies the Alg to create topographical cues as cell anchoring points to promote cell adhesion and growth. In proof-of-concept experiments, neurons exhibited robust adhesion and growth on these M-Alg scaffolds with maturation of neuronal networks evidenced by extensive neurite outgrowth and spontaneous neural activity, corroborated by the immunohistochemical detection of β-tubulin-III and synaptophysin. Future work will include mechanical characterizations of the M-Alg, systematically quantifying the effects that the density and size of the microstructured pores/channels have on regeneration of different tissues, investigations on the transplantation of M-Alg *in vivo* for tissue regeneration, longer term robustness and biodegradation of the material both *in vitro* and *in vivo*. These current studies pave the way for microstructured Alg scaffolds to serve as novel type of substrates for *in vitro* neural tissue cultivation and as guiding frameworks for nervous tissue regeneration [32].

## Supporting information

Using HCl to leach the t-ZnO components in the 3D printed scaffold.

Spontaneous neuronal activity in the established neuronal network in M-Alg scaffold.

## Acknowledgements

The authors are grateful for the financial support from the Max Planck Society.

