## Supplementary document for "Three dimensionally printed microstructured alginate scaffolds for neural tissue engineering"

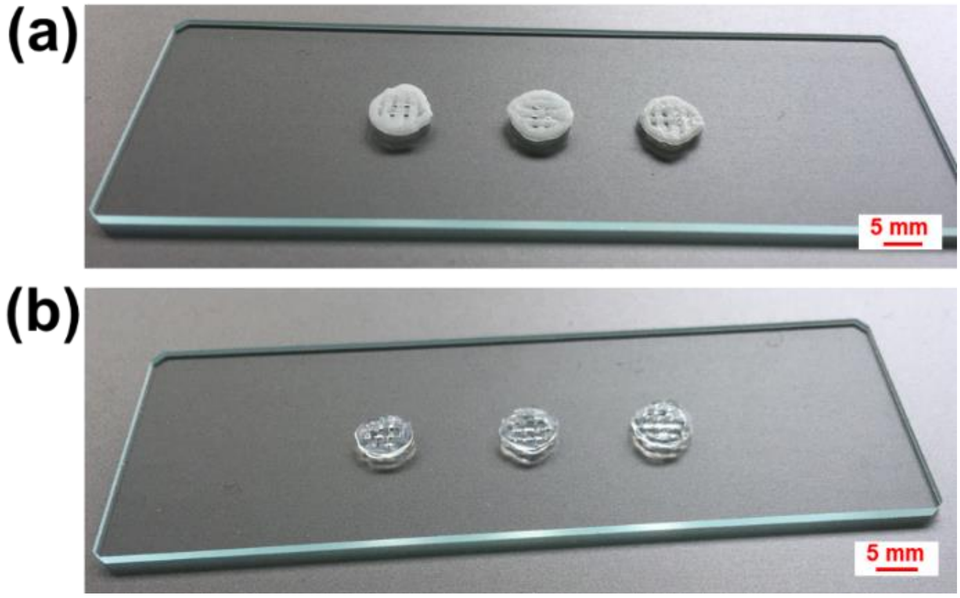


**Figure S1.** Scaffold morphology after 3D printing and t-ZnO leaching. Photos of (a) 3D printed 2%/30%/5% (w/w) Alg/Plu/t-ZnO scaffolds and (b) transparent microstructured Alg (M-Alg) scaffolds obtained after acid leaching.

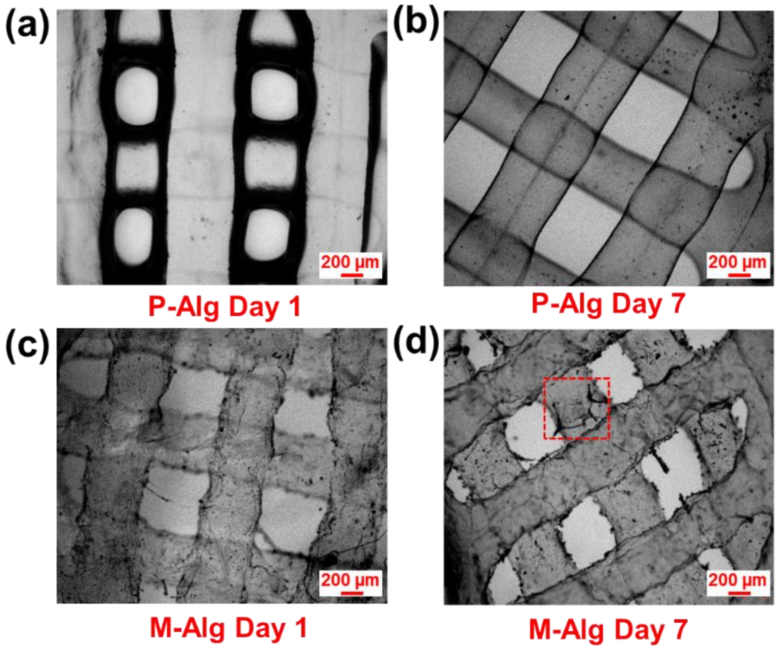


**Figure S2.** Scaffold morphology during cell culture. Optical microscope images of 3D printed (a-b) Pristine alginate (P-Alg) scaffolds and (c-d) M-Alg scaffolds on Day 1 and Day 7, correspondingly. Channels displaying tetrapod-shaped ZnO (t-ZnO) contours are highlighted with a dotted red square.

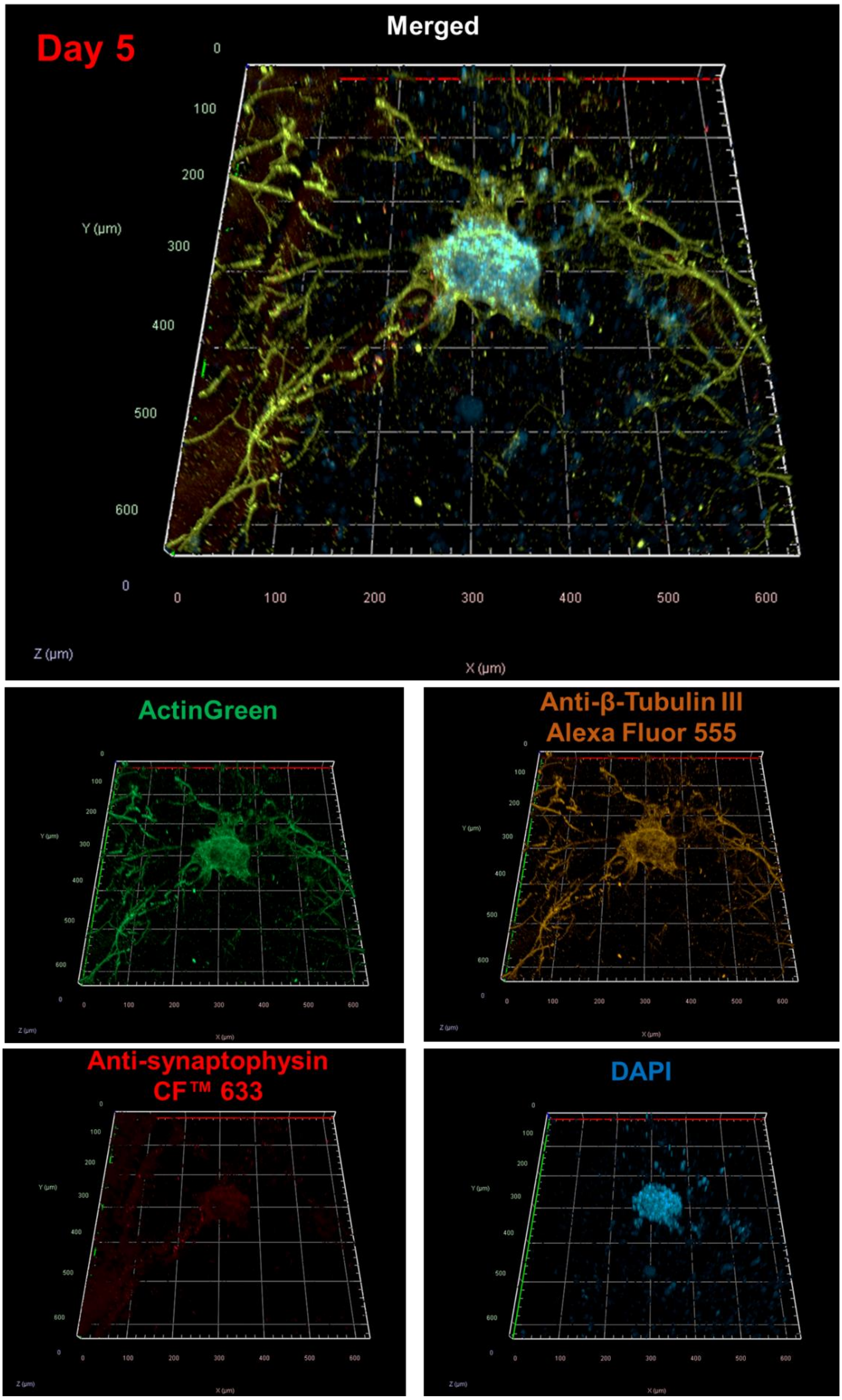


**Figure S3.** Morphology of 3D neuronal network formed within M-Alg scaffold after 5 days culture. F-actin (ActinGreen; green), β-tubulin III (Alexa Fluor 555; brown), synaptophysin ($CF^{TM}$ 633; red), nuclei (DAPI; blue).

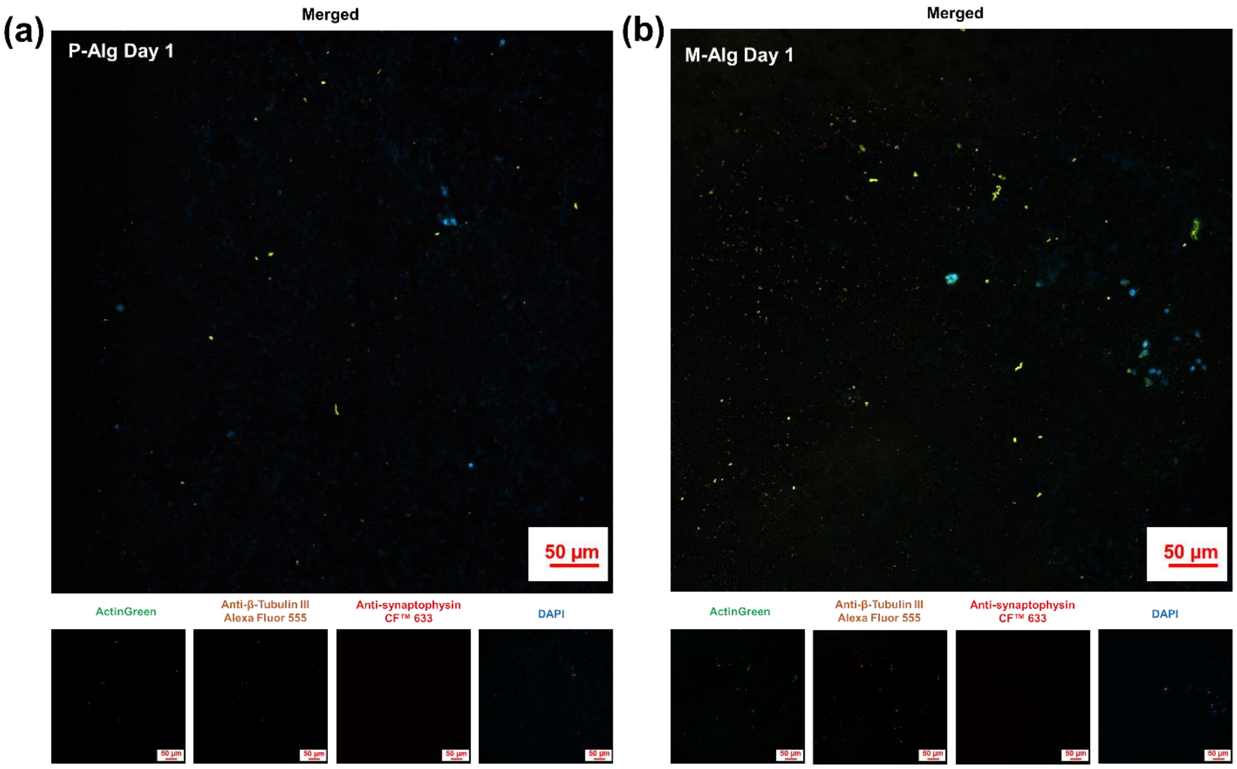


**Figure S4.** Immunofluorescence microscopy of neurons cultured on (a) P-Alg and (b) M-Alg on Day 1. F-actin (ActinGreen; green), β-tubulin III (Alexa Fluor 555; brown), synaptophysin (CF$^{TM}$ 633; red), nuclei (DAPI; blue).

**Video S1**. Using HCl to leach the t-ZnO components in the 3D printed scaffold.

**Video S2**. Spontaneous neuronal activity in the established neuronal network in M-Alg scaffold.
